# Electron transfer dictates metabolic reprogramming in proliferating cells under hypoxia

**DOI:** 10.1101/244616

**Authors:** Yuanyuan Wang, Miao Liu, Yuxia Ruan, Qiaoyun Chu, Yanfen Cui, Ceshi Chen, Guoguang Ying, Binghui Li

## Abstract

Metabolic reprogramming extensively occurs in proliferating cancer cells. This phenomenon occurs highly heterogeneously, but its origin has remained unclear. Here we use a physicochemical concept of *free electron potential* (*FEP*) and its equation of state to profile metabolites. We demonstrate that FEP change between substrates and products exactly reflects electrons dissipated in a metabolic transformation. Based on the law of conservation of electron in chemical reactions, a function of FEP change for central metabolism in proliferating cells are further derived, and it can accurately predict metabolic behaviors under hypoxia by maximizing the cellular FEP change to consume electrons. Therefore, enabling electron transfer dictates metabolic reprogramming in hypoxic cells, which underlies the major findings in cancer metabolism and is supported by our experiments. Importantly, our model established on FEP helps to reveal a combination of promising targets to inhibit tumor growth under hypoxia by blocking electron consumption, and could also guide future studies on cancer metabolism under hypoxia.

---

Although proliferating cells, such as cancer cells, develop significant metabolic heterogeneity^1–3^ to cope with the changing microenvironment, we notice that hypoxia and mitochondrial dysfunction, both of which block the electron transport chain (ETC), lead to the similar metabolic reprogramming^4,5^. It prompts us to hypothesize that the accumulation of electrons probably drives metabolic reprogramming.

As a pool of integrated chemical transformations in cells, metabolism must observe the chemical laws. If we can find out the chemical law underlying metabolic reprogramming in proliferating cancer cells, we may design a cocktail of broad-spectrum treatments for cancers regardless of the heterogeneity. Classically, chemical reactions involve the forming and breaking of chemical bonds between atoms. Since chemical bonds result from a redistribution of the outer electrons of atoms, chemical reactions may encompass electron transfer between each other or release electrons to the system. As for mammalian cells, the established fact is that oxygen is used to accept the net electrons released from the whole cellular metabolism, which is mediated by the mitochondrial electron-transport chain (ETC). Therefore, the mitochondrion plays an essential role as the electron acceptor in enabling cell proliferation in addition to energy generation^4–6^. However, when the ETC is disabled by hypoxia or pathological/pharmacological inhibition, how to enable electron transfer to support cell proliferation remains elusive. Here, we build up a physicochemical model to reveal the chemical nature of metabolic reprogramming in cells.

## The similar metabolic reprogramming induced by hypoxia and mitochondrial inhibition

Based on the observations from different research groups, hypoxia^7^ and mitochondrial deficiency^8^ led to the similar metabolic phenomena, especially the reductive carboxylation of glutamine-derived α-ketoglutarate (Figure 1A). Here we parallelably compared the metabolic effects of hypoxia and antimycin A, a mitochondrial ETC inhibitor, on cancer cells. First, we traced the metabolic pathways by using [U-^13^C-glutamine in HeLa cells. Normally, through the CAC, glutamine-derived α-ketoglutarate m+5 was predominantly oxidized to succinate m+4, malate m+4, oxaloacetate m+4 that could be converted into aspartate m+4, citrate m+4 and isocitrate m+4 (Figure 1B-1E and S1A-S1D). Upon hypoxia or ETC inhibition, glutamine-derived α-ketoglutarate m+5 was reduced by carboxylation to isocitrate m+5 and citrate m+5 that was further lysed to acetyl-CoA m+2 and oxaloacetate for the generation of aspartate m+3 and malate m+3 (Figure 1B-1E and S1A-S1D). In the meantime, glutamine-derived lactate m+2 from malate m+3 in the reductive pathway dramatically increased while that m+3 from malate m+4 in the oxidative pathway decreased under the condition of hypoxia or ETC inhibition (Figure 1F). The similar results were also obtained in 4T1 cells (Figure 1G, 1H, S1E and S1F). These results indicate that cellular metabolism is reprogrammed toward glutamine-initiated lipogenesis through the non-canonical reductive pathway (Figure 1A) in the conditions of hypoxia and ETC inhibition. In addition, hypoxia and antimycin A induced the typically increased metabolic flux of glucose to lactate (Figure S2A-S2C), and they also induced similar changes in intracellular metabolites as revealed by the metabolomics (Figure S3). Taken together, these data clearly confirm that hypoxia and ETC inhibition trigger the similar metabolic reprogramming.

**Figure 1.**
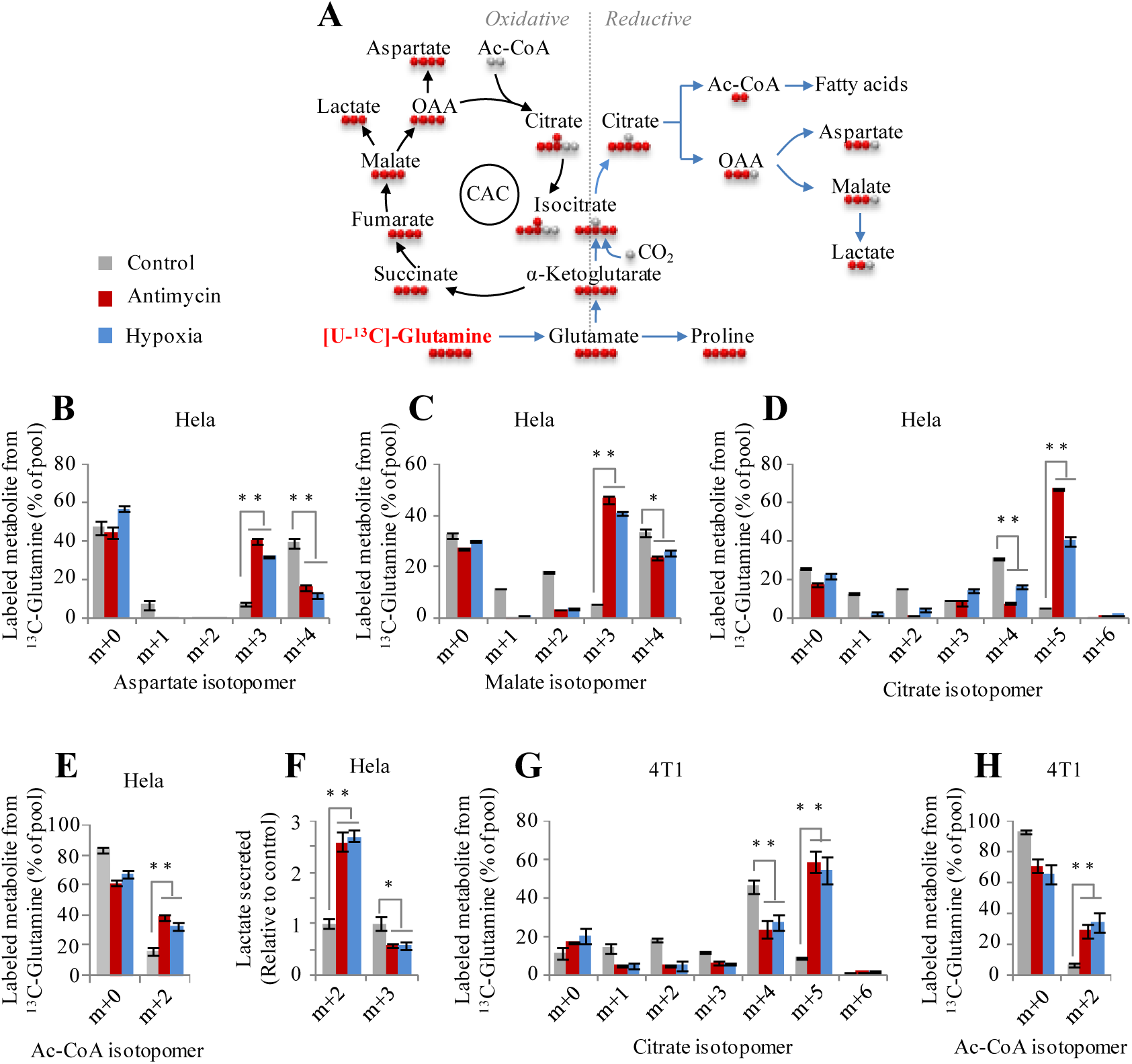
Glutamine metabolismin cancer cells under hypoxia and ETC inhibition. (a) Schematic of oxidative or reductive glutamine metabolism. (B-E) Mass isotopomer analysis of acetyl-CoA, aspartate and malate in HeLa cells cultured with ^13^C_5_-glutamine for 8h under the conditions of hypoxia or inhibition by antimycin A (1 μΜ). (F) Excretion of lactate from HeLa cells under the same condition as described in (B-E). (G and H) Mass isotopomer analysis of citrate and acteylc-CoA in 4T1 cells cultured with [U-^13^C]-glutamine under the conditions as indicated. Data are the mean ± SD for three independent cultures. \**P*<0.05; \*\**P*<0.01, Student’s *t*-test. See also Figure S1-S4.

Antimycin A can induce the accumulation of electrons. We then measured the cellular NADH and NADPH upon hypoxia and ETC inhibition. Both hypoxia and antimycin A accumulated cellular NADH and enhanced the ratio of NADH/NAD+ (Figure S4A and S4B). Although we did not observe the accumulated NADPH and increased NADPH/NADP+ ratio (Figure S4C and S4D), NADH and NADPH can be easily transhydrogenated between each other or translocated spatially by various biochemical shuttles or reversible biochemical reactions^8,9^ (Figure S5). These observations urged us to hypothesize that electron transfer may determine the metabolic reprogramming.

## The constant electron consumption between substrates and products

Electron transfer is directly associated with the oxidation and reduction of metabolites. To comprehensively understand electron transfer in cells, we derived a concept from the previous established *degree of reduction* in bioprocess engineering^10^, referred to as *free electron potential* (*FEP*)*. FEP* characterizes the potential ability of a typical biological metabolite, with the molecular formula of C_C_H_H_O_O_N_N_, to release electrons upon its complete oxidation to CO_2_, H_2_O and NH_3_ in cells. The degree of reduction of any element in a compound is equal to the valence of this element, and the degree of reduction of some elements are C = 4, H = 1, N = −3 and O = −2 in metabolites^10^. Therefore, we derived an equation of state for *FEP*,

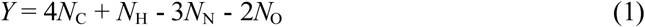

in which *Y* is used as the symbol for *FEP* and *N*_C_, *N*_H_, *N*_N_ and *N*_O_ are the number of carbon, hydrogen, oxygen and nitrogen atoms in the molecule. Here we listed *Y* values of intracellular metabolites involved in the metabolism of glucose, amino acids, nucleic acids and lipids (Table 1, and S1). Interestingly, the *Y* per carbon atom values (γ) of metabolites show somehow internal properties determined by “4” (detailed in Supplemental Discussion).

**Table 1.**
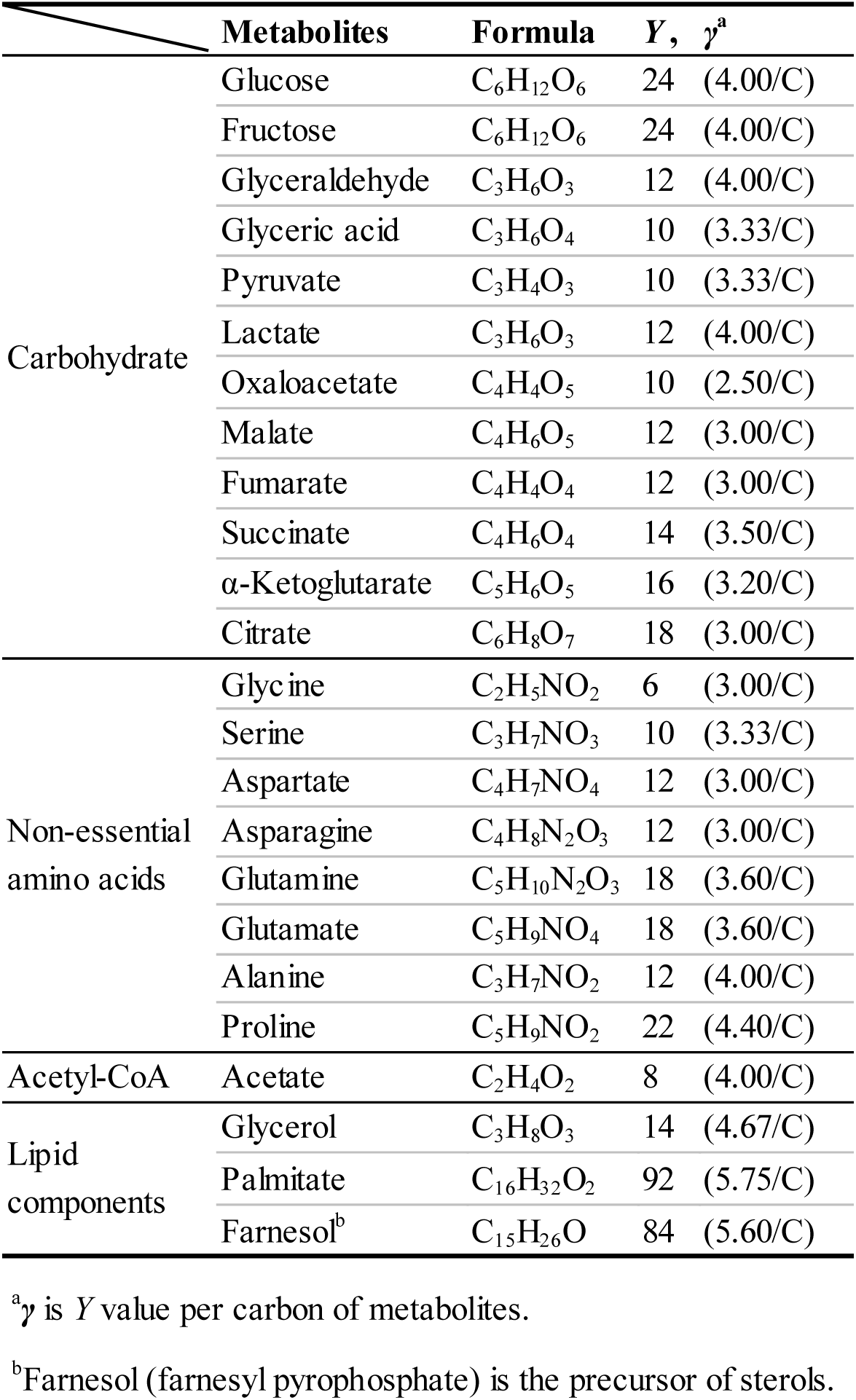
Free electron potential of metabolites

| | Metabolites | Formula | Y, $\gamma^a$ |
| --- | --- | --- | --- |
| Carbohydrate | Glucose | C<sub>6</sub>H<sub>12</sub>O<sub>6</sub> | 24 (4.00/C) |
|  | Fructose | C<sub>6</sub>H<sub>12</sub>O<sub>6</sub> | 24 (4.00/C) |
|  | Glyceraldehyde | C<sub>3</sub>H<sub>6</sub>O<sub>3</sub> | 12 (4.00/C) |
|  | Glyceric acid | C<sub>3</sub>H<sub>6</sub>O<sub>4</sub> | 10 (3.33/C) |
|  | Pyruvate | C<sub>3</sub>H<sub>4</sub>O<sub>3</sub> | 10 (3.33/C) |
|  | Lactate | C<sub>3</sub>H<sub>6</sub>O<sub>3</sub> | 12 (4.00/C) |
|  | Oxaloacetate | C<sub>4</sub>H<sub>4</sub>O<sub>5</sub> | 10 (2.50/C) |
|  | Malate | C<sub>4</sub>H<sub>6</sub>O<sub>5</sub> | 12 (3.00/C) |
|  | Fumarate | C<sub>4</sub>H<sub>4</sub>O<sub>4</sub> | 12 (3.00/C) |
|  | Succinate | C<sub>4</sub>H<sub>6</sub>O<sub>4</sub> | 14 (3.50/C) |
| | α-Ketoglutarate | C<sub>5</sub>H<sub>6</sub>O<sub>5</sub> | 16 (3.20/C) |
|  | Citrate | C<sub>6</sub>H<sub>8</sub>O<sub>7</sub> | 18 (3.00/C) |
| Non-essential amino acids | Glycine | C<sub>2</sub>H<sub>5</sub>NO<sub>2</sub> | 6 (3.00/C) |
|  | Serine | C<sub>3</sub>H<sub>7</sub>NO<sub>3</sub> | 10 (3.33/C) |
|  | Aspartate | C<sub>4</sub>H<sub>7</sub>NO<sub>4</sub> | 12 (3.00/C) |
|  | Asparagine | C<sub>4</sub>H<sub>8</sub>N<sub>2</sub>O<sub>3</sub> | 12 (3.00/C) |
|  | Glutamine | C<sub>5</sub>H<sub>10</sub>N<sub>2</sub>O<sub>3</sub> | 18 (3.60/C) |
|  | Glutamate | C<sub>5</sub>H<sub>9</sub>NO<sub>4</sub> | 18 (3.60/C) |
|  | Alanine | C<sub>3</sub>H<sub>7</sub>NO<sub>2</sub> | 12 (4.00/C) |
|  | Proline | C<sub>5</sub>H<sub>9</sub>NO<sub>2</sub> | 22 (4.40/C) |
| Acetyl-CoA | Acetate | C<sub>2</sub>H<sub>4</sub>O<sub>2</sub> | 8 (4.00/C) |
| Lipid components | Glycerol | C<sub>3</sub>H<sub>8</sub>O<sub>3</sub> | 14 (4.67/C) |
|  | Palmitate | C<sub>16</sub>H<sub>32</sub>O<sub>2</sub> | 92 (5.75/C) |
|  | Farnesol<sup>b</sup> | C<sub>15</sub>H<sub>26</sub>O | 84 (5.60/C) |

<sup>a</sup> $\gamma$ is Y value per carbon of metabolites.
<sup>b</sup>Farnesol (farnesyl pyrophosphate) is the precursor of sterols.

Equation (1) can be used to quantitatively calculate electrons required for metabolic conversions by *FEP* change, Δ*Y*_P,S_, between *Y* values of initial substrates and those of final products. As for a transformation,

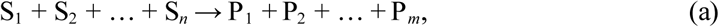

where S and P are the carbon-containing metabolites with the quantity of carbon atoms being made conserved,

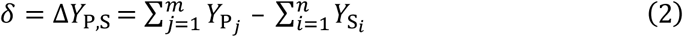

A positive *FEP* change means the consumption of electrons while a negative value indicates the release of electrons in this conversion. Phosphate group and Co-enzyme are often used to facilitate metabolic transformations, however, their addition to metabolites do not involve the transfer of electrons. Thus we only need to simply focus on the form of compounds without phosphate and Co-enzyme when Δ*Y* is calculated.

To prove equation (2), we then make the quantities of nitrogen and oxygen atoms conserved by adding *x* moles of NH_3_ and *y* moles of H_2_O as the products, and thus change reaction (a) to

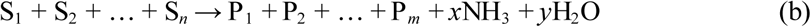

In reaction (b), the quantities of carbon, nitrogen and oxygen atoms are conserved. Notably, *x* and *y* could be negative, meaning that NH_3_ and H_2_O are actually substrates. Thus,

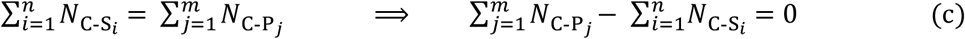

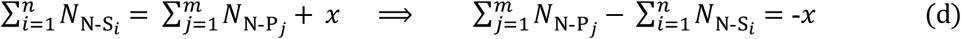

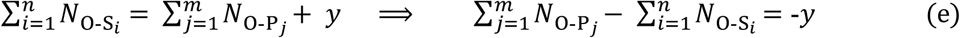

*N*_C-S_, *N*_N-S_ and *N*_O-S_ are the numbers of carbon, nitrogen and oxygen atoms in substrates while *N*_C-P_, *N*_N-P_ and *N*_O-P_ are those in products.

Based on equation (1), *Y* = 4*N_C_* + *N*_H_ - 3*N_N_* - 2*N*_O_, for metabolites,

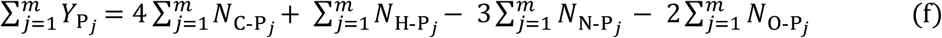

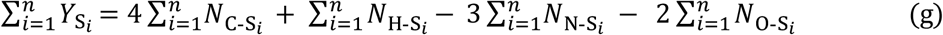

In equation (f) and equation (g), *N*_H-S_ and *N*_H-P_ are the numbers of hydrogen atoms in substrates and products.

As for reaction (b) where *Y* of NH_3_ and H_2_O is 0,

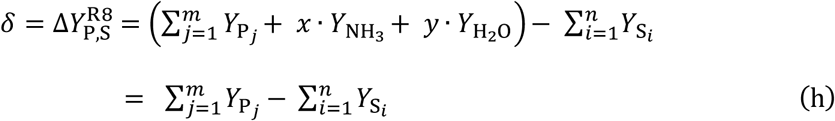

Hence

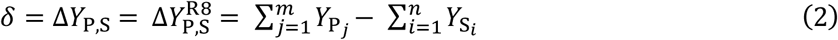

By inserting equation (f) and (g) into equation (2), we obtained

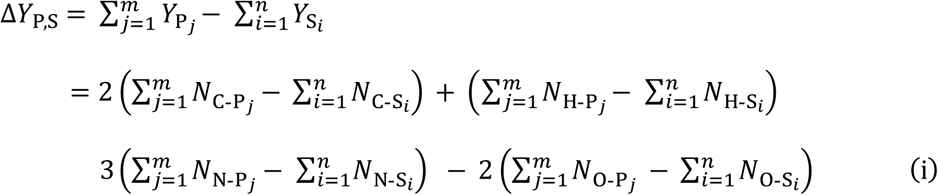

Substituting equation (c), equation (d) and equation (e) into equation (i) yields

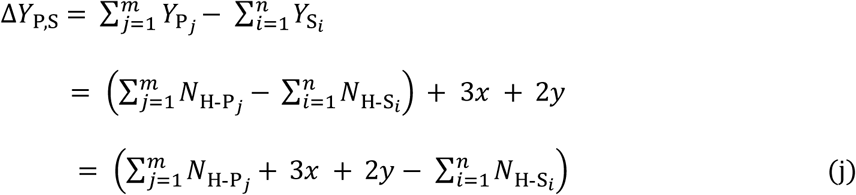

In equation (k),

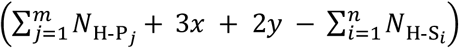

is actually the difference of the quantities of hydrogen atoms between substrates and products in reaction (b). Based on the Law of Conservation of Mass, these hydrogen atoms (the electron carriers) must be integrated into electron acceptors, such as NAD(P)^+^ and FMN/FAD, if the difference is positive. Otherwise they come from the electron donor, for example, NAD(P)H + H^+^ or FMNH_2_/FADH_2_. Therefore, Δ*Y*_P,S_ represents the number of electrons produced or consumed in reactions (a) and (b).

The conversion of substrates to products may have several metabolic routes with different energy requirements, but Δ*Y* between substrates and products keeps constant based on the above analyses.

As for most of redox reactions, NAD(P)^+^, FMN/FAD or their reductive forms are used in the electron transfer. However, in some cases, reactions may directly use other electron donors, such as tetrahydrofolic acid and ascorbic acid. The resultant oxidized electron donors, dihydrofolic acid and dehydroascorbic acid, need to be finally reduced back for the sake of redox homeostasis usually using NADPH + H^+^ as electron donor^11,12^. Therefore, in essence, these reactions equivalently consume NADPH + H^+^. In some other cases, oxygen could be directly used as electron acceptor in the reactions. Oxygen is expected to accept electrons from NADH + H^+^ through the mitochondrial electron transport chain. If oxygen accepts electrons from metabolites in reactions, the same equivalents of NADH + H^+^ are left in cells, in particular under hypoxic conditions. Thus, when used as electron acceptor, oxygen is essentially equal to NAD+. Taken together, the analyses on the transfer of global electrons in cells are not restricted to the forms of electron acceptor/donor.

## Electron transfer in proliferating cells

Proliferating cells must produce ATP and duplicate all the building bricks to make new cells. Mammalian and bacterial cells share the similar chemical compositions^13^ (Table S2). Macromolecules including proteins, nucleic acids, lipids and polysaccharides account for 87% and their precursors involving amino acids, nucleotides, fatty acids, sugars and the related intermediates make up 9.3% of cell mass. The rest chemicals are inorganic ions (3%) that are unrelated to electron transfer and other small molecules (0.7%) that could be negligible to the analysis of global intracellular electrons. Protein synthesis from amino acids, nucleic acid synthesis from nucleosides, polysaccharide synthesis from sugars and lipid synthesis from fatty acids do not involve the transfer of electrons. Therefore, here we mainly focus on the synthesis of amino acids (non-essential amino acids), nucleosides, fatty acids and sugars. The major carbon source in blood includes glucose and glutamine. We calculated the possible Δ*Y* of these building bricks, including ATP generation, lipid biosynthesis, sugar biosynthesis, amino acid synthesis and nucleotide biosynthesis, for cell proliferation based on *Y* values of metabolites in Table 1 and S1 (Figure 2A-2E and S6 and Table S3).

We can calculate the total electrons consumed by ATP generation (Figure 2A), lipid biosynthesis (Figure 2B), sugar biosynthesis (Figure 2C), amino acid synthesis (Figure 2D) and nucleotide biosynthesis (Figure 2E). Based on the law of electron conservation in chemical reactions, the total electrons produced in a cell (max*Φ*(*δ*)) should be around 0.

**Figure 2.**
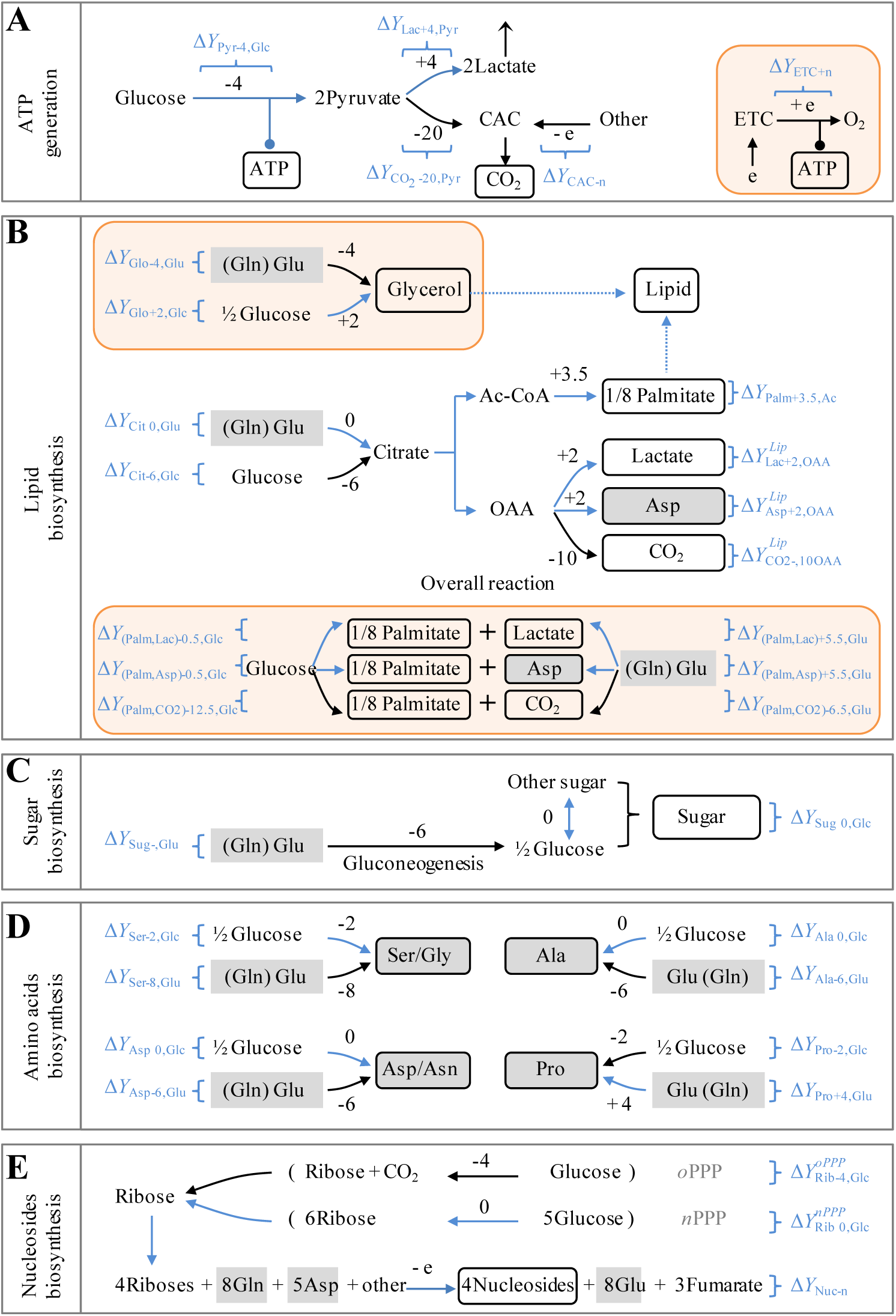
The production or dissipation of electrons (Δ*Y*) in metabolic transformations. Δ*Y* is calculated by equation (2) based on *Y* values in Table 1 and S1. (A) Δ*Y* in ATP generation. The inset is mitochondrial ATP generation. (B) Δ*Y* in the biosynthesis of lipids. Palmitate could be synthesized from glucose or glutamine. The insets are the biosynthesis of glycerol (glycerol-3-phosphate) and the overall reactions of palmitate from glucose or glutamine. (C) Δ*Y* in the biosynthesis of sugar. (D) Δ*Y* in the biosynthesis of non-essential amino acids. The carbons of amino acids could be from glucose or glutamine. (E) Δ*Υ* in the biosynthesis of nucleosides. Ribose can be generated in oxidative PPP (*o*PPP) or non-oxidative PPP (*n*PPP). The exact Δ*Y* values of ribose-initiated biosynthesis of nucleosides depend on the specified nucleoside. See Figure S6 and Table S3 for details. In all panels, numbers are Δ*Y* values in one reaction. “-” means the prodution of electrons while “+” refers to the consumption of electrons. 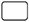 indicates the building bricks for proliferating cells. 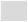 shows amino acids. Curve arrow indicates optional pathways. 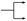 represents the coupled pathwys. Blue routes show the metabolic pathways prevailing under hypoxia. See also Figure S5 and S6.

Hence,

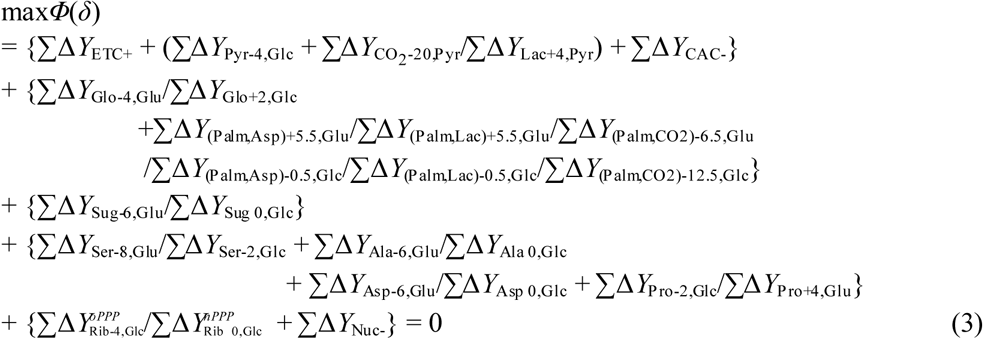

Σ means the sum of electrons from all the intracellular reactions, and variables with slashes from different metabolic pathways are optional depending on the availability of nutrients and cellular states (Figure 2). As a cell is a self-organizing system, equation (3) can be flexibly set up to 0 by using different metabolic pathways. When oxygen is sufficient, the ETC works efficiently. To support ATP generation coupled to the ETC, cells may use the metabolic pathways producing electrons.

## Metabolic reprogramming of proliferating cells upon ETC dysfunction

Under the condition of ETC dysfunction, such as hypoxia and ETC inhibition, the electron flow to oxygen is blocked (here let 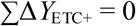), thus

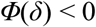

It could lead to accumulated electrons that unlikely rises infinitely (Figure S4A-S4D). Therefore, to equate *Φ*(*δ*) with 0 while still enable anabolic processes, proliferating cells have to rewire their metabolic reactions to balance electron transfer by reducing the production and enhancing the dissipation of electrons. In doing so, *Φ*(*δ*) is maximized by selecting higher Δ*Y* values from alternative metabolic pathways, thus gives

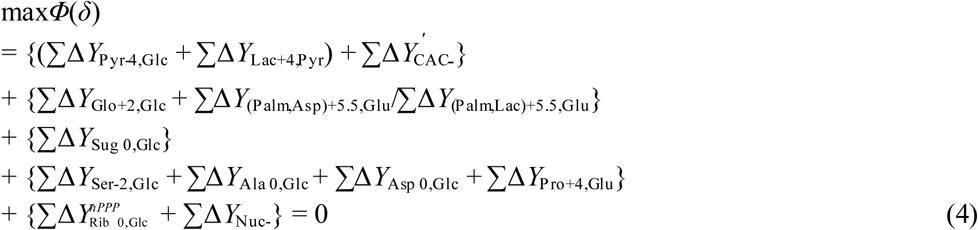

Based on max*Φ*(*δ*), glycolytic pyruvate shunts to lactate 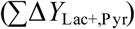 and avoids entering the CAC to produce electrons 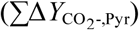, which results in the increased lactate production (Figure S2B and S2C). This pathway does not release any electron (Δ*Y* = 0), because Δ*Y*_Pyr-Glc_ is negatively equal to Δ*Y*_Lac+,Pyr_ (Figure 2A). Normally, the CAC produced a great many electrons for mitochondrial ATP generation by completely oxidizing its participants to carbon dioxides, but also provided on purpose some essential intermediates for biosynthesis. Therefore, upon ETC dysfunction, the CAC had to be shaped to a state of low activity required for cell proliferation, endowed with a 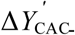 in equation (4). Our results showed that both [U-^13^C]-glucose-derived citrate m+2 and [U-^13^C]-glutamine-derived succinate m+4 were significantly reduced in the condition of hypoxia and ETC inhibition (Figure S1A, S1B and S7A-S7C), indicating a decrease in the metabolic flux of both glucose and glutamine the oxidative CAC. In addition, fatty acid biosynthesis is initiates from glutamine (Δ*Y*_(Palm,Asp)+5.5,Glu_/Δ*Y*_(Palm,Lac)+5.5,Glu_) instead of from glucose (Δ*Y*_(Palm,Asp)-0.5,Glc_/Δ*Y*(_Palm,Lac)-0.5,Glc_), and this process can dissipate electrons (Δ*Y* > 0) and concomitantly produce glutamine-derived aspartate and/or lactate through the reductive pathway (Figure 2B). These metabolic preferences explain the experimental observations^7,8^ (Figure 1 and S1-S3).

Sugar, serine, alanine and glycerol are directly derived from glucose or glycolytic intermediates, thus their biosynthesis initiated from glucose prevails in proliferating cells (Figure 3A and S8A-S8E). In contrast, proline is always synthesized from glutamate that is mainly produced from glutamine (Figure 3B and S8F). Consistently, these metabolic preferences support max*Φ*(*δ*), but hypoxia and ECT inhibition promote glucose-derived glycerol 3-phosphate (Figure 3A) and glutamine-derived proline (Figure 3B) because of their positive Δ*Y* values (Δ*Y*_Glo+2,Glc_ and Δ*Y*_Pro+4,Glu_ > 0) (Figure 2C and 2D), which contributes to max*Φ*(*δ*).

**Figure 3.**
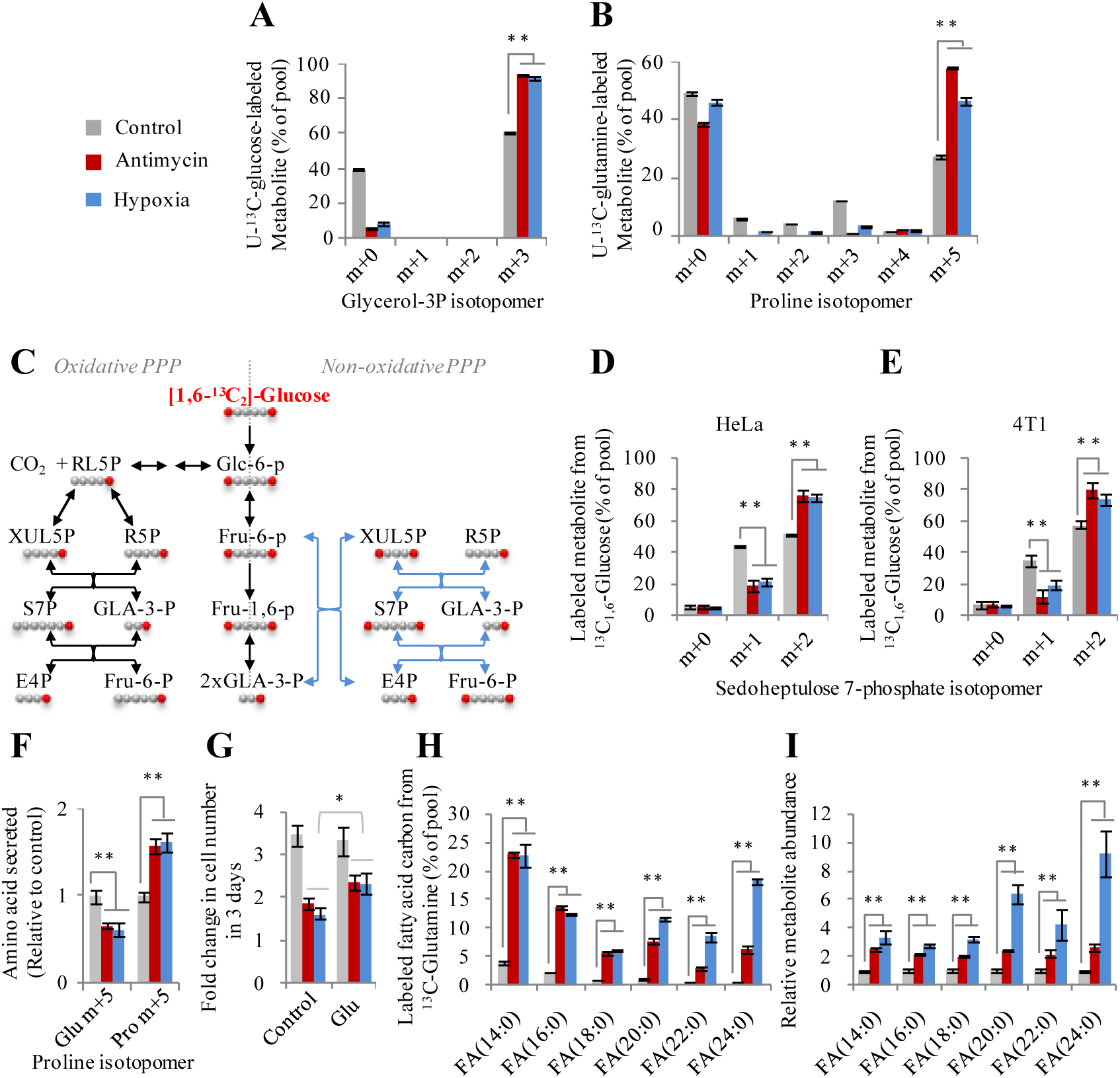
Metabolic reprogramming in cancer cells according to max*Φ*(*δ*). (A) Mass isotopomer analysis of glycerol 3-phosphate in HeLa cells cultured with ^13^C_6_-glucose for 8h under the conditions of hypoxia or inhibition by antimycin A (1 μM). (B) Mass isotopomer analysis of proline in HeLa cells cultured with ^13^C_5_-glutamine for 8h under the conditions of hypoxia or inhibition by antimycin A (1 μM). (C) Schematic for oxidative pentose phosphate pathway (oPPP) and non-oxidative pentose phosphate pathway (nPPP). (D and E) Mass isotopomer analysis of sedoheptulse-7 phosphate in HeLa and 4T1 cells cultured with [1,6–^13^C_2_]c-glucose under the conditions as indicated. (F) Excretion of glutamate and proline from HeLa cells cultured with ^13^C_5_-glutamine for 8h under the conditions of hypoxia or inhibition by antimycin A (1 μM). (G) Effect of glutamate on cell growth. HeLa cells were cultured with or withour 10 mM glutamate in the condition of hypoxia or inhibition by antimycin A (1 μM) for 3 days. (H) Labeled carbons in fatty acids in HeLa cells cultured with ^13^C_5_-glutamine for 48h under the conditions of hypoxia or inhibition by antimycin A (1 μM). (I) Relative abundance of fatty acids in HeLa cells cultured with ^13^C_5_-glutamine for 48h under the conditions of hypoxia or inhibition by antimycin A (1 μM). Data are the mean ± SD for three independent cultures. \**P*<0.05; \*\**P*<0.01, Student’s *t*-test. See also Figure S7-S12.

There are currently controversial opinions on the roles of NADPH produced in the oxidative pentose phosphate pathway (PPP) in cancers. Supporting data mainly focus on the antioxidant activity of PPP-produced NADPH^14^. However, many reports recently showed that the oxidative PPP could be shut down in cancer cells^15–17^. Moreover, the epidemiological investigations demonstrated that G6PD deficiency, the rate-limiting enzyme in the oxidative pathway, did not affect the incidence of cancers^18,19^. As shown in equation (4), max*Φ*(*δ*) prefers the non-oxidative PPP 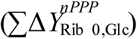 to reduce the production of electrons in the condition of hypoxia and ECT inhibition (Figure 2E). Our result also confirm that the fraction of [3–^2^H]-glucose-derived [^2^H]-labeled NADPH m+1 via the oxidative PPP was observed to decline upon hypoxia and ETC inhibition (Figure S9A and S9B). The analysis of enrichment of [1,6–^13^C_2_]-glucose in sedoheptulose 7-phosphate, a specific intermediate of PPP, further demonstrated that sedoheptulose 7-phosphate m+2 in the non-oxidative PPP increased while sedoheptulose 7-phosphate m+1 via the oxidative PPP decreased under hypoxia or ETC inhibition (Figure 3C-3E). These data suggest that the non-oxidative PPP, instead of the oxidative PPP, is preferentially used by cancer cells under hypoxia.

Furthermore, simplifying equation (4) by removing all the variables with Δ*Y* = 0 and rearranging it yielded

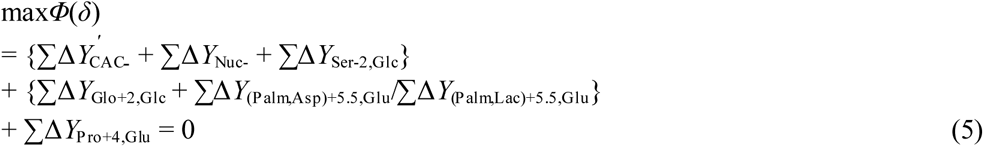

Electrons were minimally produced by the anabolic transformations including CAC, serine synthesis and nucleic acid synthesis, here designated as ΣΔ*Y*_-_. In fact, cells may release additional electrons (detailed in Supplemental Discussion). The assimilation of glycerol is coupled to glutamate-initiated lipid synthesis (Figure 2B), with the electron production being denoted as ΣΔ*Y*_Lip+,Glu_. Thus, equation (5) is simplified as equation (6),

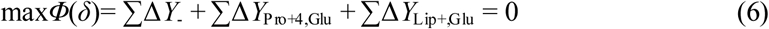

Obviously, glutamine/glutamate-initiated syntheses of both lipid and proline essentially play a determinative role in metabolically dissipating electrons to support cell proliferation under the condition of hypoxia and ETC inhibition.

Glutamate is the product of glutamine degradation and can be directly excreted if beyond the cellular requirement. Therefore, we measured the level of glutamate and proline in the medium, and found that glutamine-derived glutamate reduced while glutamine-derived proline significantly enhanced in the medium under hypoxia or ETC inhibition (Figure 3F). These results suggest that the metabolic conversation of glutamate to proline is advantageous to cell survival under hypoxia or ETC inhibition. This speculation was further confirmed by the result that supplementation with glutamate facilitated cell growth under hypoxia or ETC inhibition (Figure 3G).

Furthermore, we demonstrated that glutamine-^13^C was significantly enriched almost in all the measured fatty acids upon hypoxia and ETC inhibition (Figure 3H and S10A-S10I). Moreover, the contents of these fatty acids were boosted under these conditions (Figure 3I and S11), suggesting that cancer cells can adapt to hypoxic microenvironments by accumulating lipid mass. This finding could explain why cancer cells contain increased numbers of lipid droplets^20^.

## The metabolic equivalent of glutamine-associated metabolic reactions

As revealed in equation (6), cancer cells can accommodate to hypoxia by initiating glutamine/glutamate-derived lipid and/or proline. Here the overall Δ*Y* value for the metabolic equivalent of each glutamine/glutamate-associated metabolic reactions (GMR) is denoted as

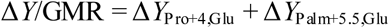

where palmitate is used for quatitation. The total available glutamate (Glu_t_) could be primarily diverted to synthesize lipid to consume more electrons under hypoxia (Figure 4A). Hence,

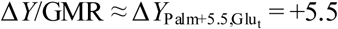

**Figure 4.**
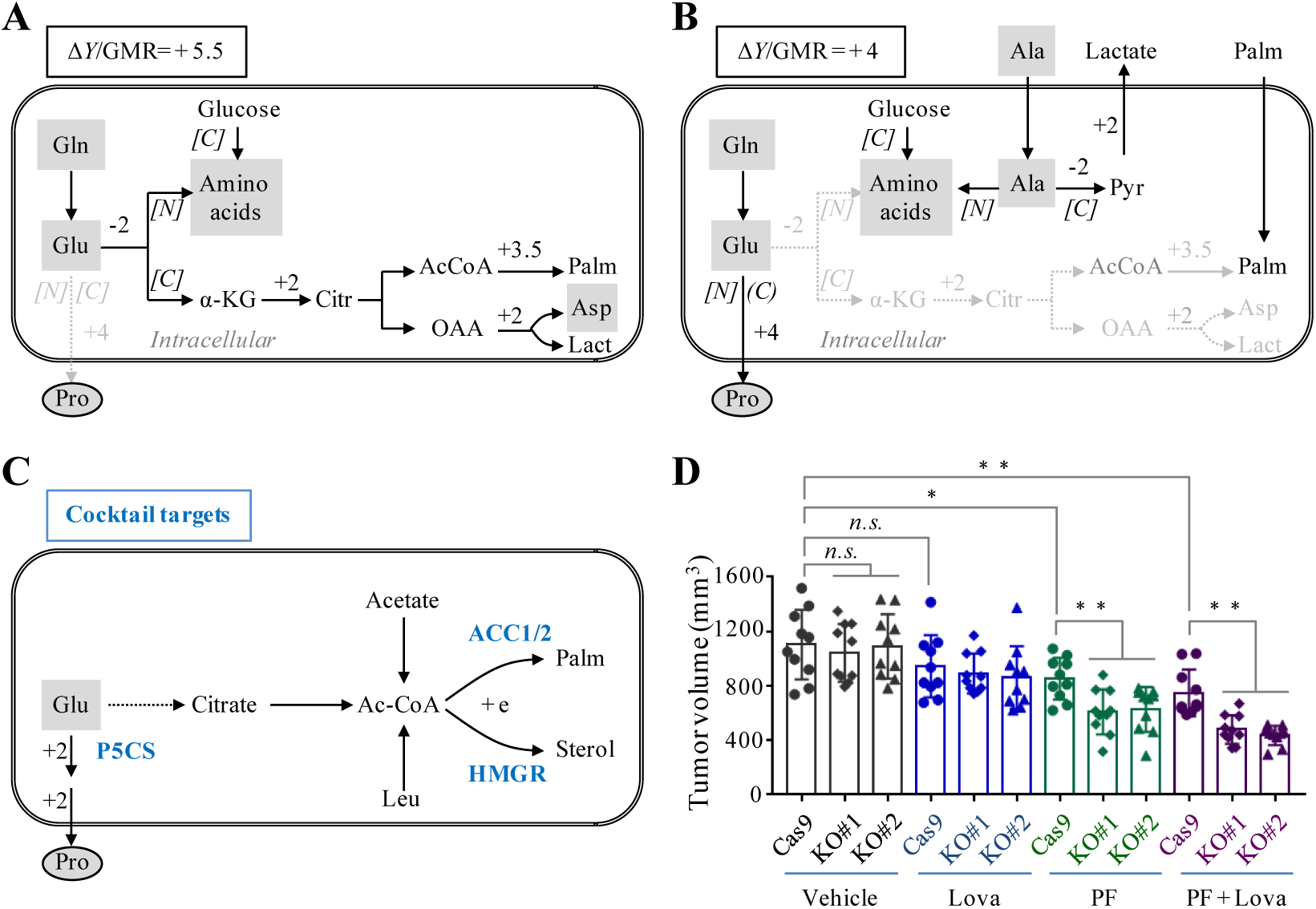
Potential metabolic adaption in proliferating cells with ETC dysfunction. ΔΥ/GMR referred to the overall ΔΥ value for the equivalent of each glutamate-associated metabolic reactions. (A) A model to show the glutamate-initiated palmitate biosynthesis. (B) A model to show glutamate-derived secretory proline. Cells use exogenous lipids/palmitates, and glutamate is used to synthesize proline that is secreted out of cells. Amino acids are absorbed or assumed to synthesize from alanine amino group. (C) Potential targets to block electrons-consuming metabolic pathways. (D) Simultaneous inhibition of P5CS, ACC1/2 and HMGR synergistically suppress tumor growth *in vivo*. P5CS, pyrroline-5-carboxylate synthase, is the key enzyme in the synthesis of proline, and it is knockouted in 4T1 cells. PF-05175157 (PF) and lovastatin (Lova) are the inhibitors of ACC1/2, acetyl-CoA carboxylase, and HMGR, 3-hydroxy-3-methyl-glutaryl-coenzyme A reductase. Here tumors from vehicle-treated and 30 mg/kg Lova plus 100 mg/kg PF-treated nu/nu mice on day 16. \**P*<0.05, Student’s *t*-test (n=10). See also Figure S13 and S14.

Nevertheless, the conversion of glutamate to proline has a more efficient Δ*Y*/ATP ratio of +4, compared to the glutamate-initiated palmitate synthesis with a Δ*Y*/ATP of +2.93 (Figure S12). Therefore, cells may utilize extracellular lipid and shift glutamate from lipid synthesis to secretory proline synthesis in the hypoxic *in vivo* microenvironment, only if they can directly acquire amino acids or obtain nitrogen source for amino acid synthesis, such as alanine, the second most abundant amino acid in human blood (Figure 4B). The carbon atoms of alanine can be converted to excretory lactate, and this process does not produce electron (Δ*Y* = 0) (Figure 4B). As for these transformations,

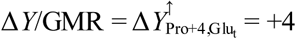

where 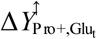 is the Δ*Y* of secretory proline synthesized from the total available glutamate (Figure 4B).

In addition, cells could synthesize lipid from other carbon sources with positive Δ*Y*/GMR. Since acetyl-CoA is the precursor of fatty acids, Δ*Y*/GMR depends on the synthesis of acetyl-CoA from other nutrients. Hence,

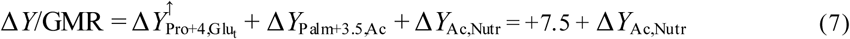

As listed in Table S4, some nutrients have a positive Δ*Y*/GMR. In particular, acetate with Δ*Y*/GMR = +7.5 (Table S4 and Figure S13A) is higher than that of glutamine/glutamate with Δ*Y*/GMR = +4 ~ +5.5, and thus could be preferentially utilized for lipogensis under hypoxia^21,22^. In addition, leucine with Δ*Y*/GMR = +5.5 (Table S4 and Figure S13B) should also be an alternative carbon source for lipid biosynthesis. These metabolic pathways could be cell type-specific and context-dependent, but all of them are centered on biosynthesis of lipid and proline.

## The potential treatments for cancers

Based on all the above analyses, the simultaneous blockage of both proline and lipid syntheses should effectively disable *in vivo* growth of cancers due to their hypoxic micro-environment. P5CS is the critical enzyme involved the proline biosynthesis pathway, and ACC1/2 and HMGR are the rate-limiting enzymes in the biosynthesis of lipids, mainly fatty acids and sterols^23,24^ (Figure 4C). Since no inhibitor against proline synthesis is available, we generated P5CS-null and -wild type HeLa and 4T1 cells. However, inhibition of proline synthesis by depleting P5CS marginally sensitized cells to hypoxia in the proline-contained medium (Figure S13C and S13D). This should be attributable to the fact that cells usually have active *de novo* lipogenesis, the major electron-dissipated process. Consistently, P5CS knockout did not affect tumor growth but sensitize tumors to PF-05175157 and/or lovastatin, the inhibitors of ^25^ACC1/2 and ^26^HMGR (Figure 4D and S14), suggesting that praline biosynthesis can partially compensate the blockage of lipogenesis. Therefore, inhibitions of the biosynthesis of proline and lipid can synergistically suppressed tumor growth.

## Discussion

Due to the fast growth rate and poor vasculature, tumor cells often suffer from insufficient supply with nutrition and oxygen^3^. To maintain uptake of nutrients, tumor cells usually enhance expression of their transporters^27–29^. However, oxygen passively diffuses into cells, and thus tumor cells are unable to acquire enough oxygen under hypoxia. Instead, they adapt to such harsh conditions by genetically or epigenetically optimizing their metabolism according to max*Φ*(*δ*), as revealed in the current study. Therefore, cancer cells still display some hypoxia-associated metabolic phenomena even if they are *in vitro* cultured. This essentially interprets metabolic reprogramming in cancers^1,2,30^, such as abnormal glucose metabolism^31^, ectopic utilization of alanine^32^ and branched-chain amino acid^33,34^ and *de novo* lipid synthesis from glutamine^7,8,35^ or acetate^21,22,36,37^.

In addition to the interpretations on the typical metabolic phenomena in cancers, max*Φ*(*δ*) chemically explains why hypoxia and ETC inhibition almost induce the same metabolic reprogramming, reveals that hypoxia and ETC inhibition stimulate the biosynthesis of glucose-derived glycerol 3-phosphate and glutamine-derived proline, and emphasizes the role of non-oxidative PPP, proline excretion and lipid droplets in cancer cells under the hypoxic conditions. Complementary to several recent reports showing that proline synthesis may play a critical role in supporting *in vivo* growth of some cancers^38, 39^, our model reveals that the process of proline biosynthesis, rather than proline itself, is important for cancer cell growth under hypoxic conditions. Catabolism of branched-chain amino acids, including leucine, isoleucine and valine, share the first two enzymes, branched-chain amino acid transaminase (BCAT) and branched-chain α-keto acid dehydrogenase complex (BCKD), thus the three amino acids are often equally studied for caners^33,34^. However, our theory distinguishes them by their metabolic equivalents, Δ*Υ*/GMR values (Table S4), and indicates that only leucine could be potentially advantageous to cell survival under hypoxia. Although this speculation needs further experimental validation, leucine indeed shows some specific functions, such as regulation of mTOR pathway^40^, different from isoleucine and valine. It is impossible to cover all the metabolic transformations in the current study, but equation (2) can be conveniently used to analyze the additional metabolic pathways. To maximize *Φ*(*δ*), any metabolic transformation with Δ*Υ* > 0 could be of benefit for the survival of cancer cells under hypoxia or with deficient mitochondria. Furthermore, equation (2) in combination with its derivative equation (7) can calculate the metabolic equivalent of nutrients to determine its potential assimilation into lipids.

Metabolic reprogramming can be promoted by numerous signal transduction pathways with a high degree of heterogeneity in cancer cells^1–3^. Importantly, bypassing these intricate and interchangeable cascades, now we can mainly control electron transfer under hypoxia by inhibiting the synthesis of both proline and lipid through a cocktail of treatments. Moreover, these inhibitors are expected to be nontoxic to normal cells that can survive on blood lipid and proline supply. Therefore, they appear to be a promising broad-spectrum treatment for solid tumors, and can be potentiated by disabling max*Φ*(*δ*). One may expect some strategies for counteracting max*Φ*(*δ*), inhibiting lactate dehydrogenases to increase electron production from intracellularly metabolized pyruvate, blocking angiogenesis to reduce supply with oxygen as the electron acceptor, impairing the ETC with biguanides to exacerbate hypoxia, starving with serine/glycine and nucleotides to force cells synthesize these metabolites (Δ*Y* < 0), and so on. Generally, any means to neutralize max*Φ*(*δ*) established on equation (2) could be a potential treatment for cancers alone or in combination with inhibition biosynthesis of lipid and proline. (See Supplemental Discussion for more)

**Online Contents** including Supplemental Figure S1-S14, Table S1-S4, Methods, Discussion.

## Authors Contributions

B.L. conceived and designed the study, and developed the model and derived the equations; Y.W. and M.L. performed experiments; Y.W., M.L. and B.L. analyzed the data; Y.R., Q.C. and Y.C. created some constructs and cell lines; C.C. provided constructive advice; G.Y. provided constructive advice and in particular coined the term of "*free electron potential*" whose symbol, "*Y*", is the first letter of his last name; B.L. wrote the paper.

## Acknowledgements

We thank Dr. Qunying Lei (Fudan University, China) and Dr. Wei Du (University of Chicago, USA) for carefully reading through the manuscript and valuable discussion and also thank Dr. Xiaohui Liu (Metabolomics Facility at Tsinghua University Branch of China National Center for Protein Sciences, China) for technical help. This work is supported by Grants 81622037, 81372185 and 81672762 from Natural Science Foundation of China.

